# Psilocybin Reduces Grooming in the SAPAP3 Knockout Mouse Model of Compulsive Behaviour

**DOI:** 10.1101/2024.10.23.619763

**Authors:** James J Gattuso, Carey Wilson, Anthony J Hannan, Thibault Renoir

## Abstract

Psilocybin is a serotonergic psychedelic compound which shows promise for treating compulsive behaviours. This is particularly pertinent as compulsive disorders require research into new pharmacological treatment options as the current frontline treatments such as selective serotonin reuptake inhibitors, require chronic administration, have significant side effects, and leave almost half of the clinical population refractory to treatment.

In this study, we investigated psilocybin administration in male and female SAPAP3 knockout (KO) mice, a well-validated mouse model of obsessive compulsive and related disorders. We assessed the effects of acute psilocybin (1 mg/kg, intraperitoneal) administration on head twitch and locomotor behaviour as well as anxiety- and compulsive-like behaviours at multiple time-points (1-, 3- and 8-days post-injection).

While psilocybin did not have any effect on anxiety-like behaviours, we revealed for the first time that acute psilocybin administration led to enduring reductions in compulsive behaviour in male SAPAP3 KO mice and reduced grooming behaviour in female WT and SAPAP3 KO mice. We also found that psilocybin increased locomotion in wild-type littermates but not in SAPAP3 KO mice, suggesting *in vivo* serotonergic dysfunctions in KO animals. On the other hand, the typical head-twitch response following acute psilocybin (confirming its hallucinogenic-like effect at this dose) was observed in both genotypes.

Our novel findings suggest that acute psilocybin may have potential to reduce compulsive-like behaviours (up to 1 week after a single injection). Our study can inform future research directions as well as supporting the utility of psilocybin as a novel treatment option for compulsive disorders.

## Introduction

Obsessive-compulsive disorder (OCD) is a debilitating mental health condition characterized by persistent, intrusive thoughts (obsessions) and repetitive behaviours or mental acts (compulsions) performed in response to these thoughts (Goodman et al., 2014). The disorder typically manifests during adolescence or early adulthood and can significantly impair daily functioning (Abramowitz et al., 2009). Common obsessions encompass themes of contamination, harm, and perfectionism, while compulsions often involve rituals such as washing, checking, or counting (Veale & Roberts, 2014). Individuals with OCD often experience distress and anxiety related to their obsessive thoughts, which can cause compulsive behaviours as a means of alleviating the distress (Abramowitz et al., 2009; Goodman et al., 2014; Veale & Roberts, 2014). OCD affects a substantial portion of the global population, with a lifetime prevalence estimated at around 1-3% (Goodman et al., 2014). Both children and adults can be affected, with a somewhat higher prevalence in females (Fawcett et al., 2020).

The aetiology of OCD is multifactorial, involving a complex interplay of genetic, neurobiological, and environmental factors. Family studies suggest a heritable component, with certain genes implicated in susceptibility (Stewart et al., 2013). Neurotransmitter signalling dysregulation, particularly involving serotonin, is also implicated in the pathophysiology of OCD (Baumgarten & Grozdanovic, 1998; Westenberg et al., 2007). Environmental factors, such as those associated with childhood trauma or stress, may contribute to the development or exacerbation of symptoms (Wilson et al., 2023).

SAPAP3 (also known as DLGAP3) is a protein that plays a crucial role in the function and organization of synapses in the brain. It is a member of the PSD-95/Discs Large/ZO-1 (PDZ) domain-containing proteins family and is primarily expressed in the central nervous system (Welch et al., 2004). SAPAP3 is localized in the postsynaptic density of excitatory synapses and is involved in the regulation of synaptic signalling and plasticity (Chen et al., 2011; Lousada et al., 2021; Welch et al., 2004, 2007). Additionally, allelic variations in the human orthologue of SAPAP3 has been implicated in OCD in clinical genetic studies (Bienvenu et al., 2009; Naaz et al., 2022; Stewart et al., 2013; Züchner et al., 2009).

To better understand the functional significance of SAPAP3, researchers have genetically engineered SAPAP3 knockout (KO) mice, which lack the expression of the SAPAP3 gene. Welch et al. (2007) demonstrated that SAPAP3 KO mice displayed deficits in certain forms of synaptic plasticity and showed abnormal behaviours related to anxiety and repetitive and compulsive behaviours (i.e., excessive grooming), resembling some features of neuropsychiatric disorders like obsessive-compulsive disorder (OCD). Notably, the excessive grooming behaviour in SAPAP3 KO mice can lead to face and body lesions indicating that this is a compulsive-like behaviour, because despite causing self-harm the mice cannot reduce the behaviour (Welch et al. 2007).

More recent evidence shows that, at the behavioural level, SAPAP3 KO mice also exhibit hypolocomotion, reduced body weight, tic-like head/body shakes, disrupted habit formation, impaired reversal learning, altered pre-pulse inhibition and valance processing (Gattuso et al., 2023a; Hadjas et al., 2019; Kajs et al., 2022; Lamothe et al., 2023; Manning et al., 2019, 2021). At the molecular level, SAPAP3 KO mice display serotonergic and glutamatergic dysfunction, altered cortical-striatal circuitry, reduced axon calibre, mGlu5-dependent silencing of AMPA receptor synapses, and lateral orbitofrontal dysfunction (Hadjas et al., 2020; Lei et al., 2019; Lousada et al., 2021; Ramírez-Armenta et al., 2022; Wan et al., 2011; Wood et al., 2018). Furthermore, recent evidence has found that rescue of the SAPAP3 gene in striatal astrocytes and normalising striatal astrocyte morphology and function reduced the anxiety-like and compulsive-like phenotype in SAPAP3 KO mice (Soto et al., 2023, 2024). Since SAPAP3 KO mice were developed by Welch et al. (2007) they have been established as one of the most well-validated mouse models of OCD (Ahmari, 2016; Wilson et al., 2023) and obsessive-compulsive related disorders (OCRD) (Lamothe et al., 2023).

The selective serotonin reuptake inhibitor (SSRI) fluoxetine (a front-line pharmacological treatment for clinical OCD) reduces the severity of the excessive grooming and anxiety-like behaviour observed in SAPAP3 KO mice (Welch et al., 2007). However, fluoxetine remains an incomplete treatment for OCD as it requires chronic administration and months to reduce symptomology, leaves a substantial number of individuals refractory to treatment (around 30 – 40%), has a considerable side-effect profile and can lead to withdrawal symptoms upon discontinuation (Association et al., 2007; Dougherty et al., 2004; Pittenger & Bloch, 2014; Renoir, 2013). Therefore, identifying a fast and effective pharmacological treatment will reduce OCD morbidity.

Psilocybin is a psychedelic prodrug that, upon entering the body, rapidly dephosphorylates to its active drug form psilocin which is a partial agonist of 5-HT_1A_ (Ki ∼ 567.4 nM), 5-HT_2A_ (Ki ∼ 107.2 nM) and 5-HT_2C_ (Ki ∼ 97.3 nM) receptors (Besnard et al., 2012; Halberstadt et al., 2011; Nichols, 2016) and has shown promise for the treatment of a range of neuropsychiatric disorders, namely depression and anxiety (Gattuso et al., 2023b). It has been observed that the plasma elimination half-life of psilocin is approximately 2 hours in humans and 0.9 hours in mice (Thomann et al., 2024).

In the context of OCD, psilocybin has also been able to reduce marble-burying behaviour in rodents and reduced obsessive-compulsive symptomatology in humans (Matsushima et al., 2009; Moreno et al., 2006; Odland et al., 2021; Singh et al., 2023). However, preclinical trials assessing psilocybin on compulsive-like behaviour rely heavily on the marble-burying test, which has been reported to have questionable reliability and validity due to a lack of standardised parameters between laboratories (Wilson et al., 2023). Additionally, in a study where psilocybin reduced symptomatology in patients with OCD (Moreno et al., 2006), the study contained only nine patients, lacked a placebo control, contained only self-reported measures and long-term effects were not assessed. Thus, as informed by previous literature reviews (Gattuso et al., 2023b; Pedicini & Cordner, 2023) we are investigating whether psilocybin will reduce anxiety and compulsive behaviour in a well-validated mouse model (SAPAP3 KO mice). We found that psilocybin did not alter anxiety-like behaviour but led to enduring reductions in compulsive behaviour in SAPAP3 KO mice. These findings can further elucidate the treatment potential of psilocybin, and inform future preclinical and clinical research into psilocybin treatment for OCD and OCRD.

## Materials and Methods

### Animals

Male and female SAPAP3 KO mice and wild-type (WT) littermates were bred from heterozygous x heterozygous breeding pairs on a C57BL/6J background and were derived from a colony initially established at the Florey Institute of Neuroscience and Mental Health. Mice were housed in groups of 2–5 same sex/genotype mice in individually ventilated cages until weaning at four weeks of age, when they were then transferred to standard open-top boxes (34 × 16 × 16 cm; 2-5 mice/box). Mice had *ad libitum* access to standard chow and drinking water, in the 12:12 light/dark cycle (lights were on from 7am – 7pm). Mice were 5– 7 months of age at the time the experiments were performed. In total 55 male mice were used (27 WT and 28 KO) and 36 female mice were used (13 WT and 23 KO). The same mice were used across behavioural experiments. All animal experiments were approved by the Florey Institute Animal Ethics Committee, following the guidelines of the National Health and Medical Research Council (NHMRC). KO mice were checked weekly for lesions and their nails were cut if upon visual inspection a body and face lesion was beginning to develop. Nails were cut at least 48 hours before any behavioural experiment. For the male SAPAP3 KO mice 15/28 mice developed lesions. For the female SAPAP3 KO mice, 4/23 developed lesions.

### Drugs

Psilocybin (1 mg/kg), provided by Usona Institute Inc., was dissolved in 0.9% saline solution, and administered via intraperitoneal (i.p.) injection. This dose was based on previous research (Hesselgrave et al., 2021; Shao et al., 2021) where it yielded hallucinogenic-like properties (by eliciting a significant number of head twitches) and exhibited therapeutic behavioural effects as well as increasing dendritic growth in the prefrontal cortex (Shao et al., 2021). Control mice received vehicle treatment under the same conditions (volume of injection 10 ml/kg, 0.9% saline, i.p.).

### Behavioural Experiments

All behavioural testing was conducted during the light cycle (9 am - 4 pm). In all experiments, mice were acclimatised to their environment for at least 30 minutes before testing. All experiments were performed with the experimenter blinded to experimental conditions. Based on the light-dark box and grooming baseline measurements (1-week prior to drug administration), mice were pseudo-randomised to their treatment condition to ensure no treatment differences between mice were present prior to drug administration. Behavioural Experiments were conducted from April 2023 – October 2023.

#### Locomotor Activity

Locomotor activity was measured for 15 minutes before and 60 minutes after acute psilocybin administration by placing mice in a 26 x 26 x 38 cm photo-beam activity chamber (Coulbourn Instruments, PA, USA) (Wilson et al., 2020). In our analysis, we analysed distance travelled (cm) via 5-minute bins. We used the 10-15-minute bin before injection as our baseline (time point 0).

#### Head-Twitch Response (HTR)

In the 15 minutes immediately after injection, mice were placed in the locomotor cell and were video recorded from a top view (the video data was captured at a resolution of 1920 × 1088 pixels with a frame rate of 25 frames per second). The behaviour was manually scored by a blinded and trained researcher. The HTR was measured while mice were completing their locomotor behaviour.

#### Light-Dark Box (LDB)

The LDB apparatus consisted of square photo-beam activity arenas (26 × 26 × 38 cm) that are divided into light and dark compartments with a black Plexiglas insert. The light compartment was brightly lit at 750 ± 10 lux. Following a protocol previously published (Gattuso et al., 2023a), mice were placed in the dark compartment, through a lid on the top of the insert, and then allowed to explore the arena for 10 minutes. Anxiety-like behaviour was assessed at three time-points (i.e., Baseline, 24 hours, 72 hours, and 8 days after acute psilocybin injection) by measuring the percentage of time spent in the light compartment of the LDB. Additionally, subsidiary measures such as the distance travelled and the number of transitions between the light and dark compartment were also calculated.

#### Grooming

Grooming behaviours were assessed at 4 timepoints (i.e., baseline, 24 hours, 72 hours and 8 days after psilocybin) following the LDB (on the same day, after habituating to the environment). For instance, mice were habituated to the LDB room (30 minutes, in home cage) then conducted the LDB test, were then transferred to the grooming behavioural room (in their home cage) and habituated for 30 minutes before grooming was assessed. Mice were individually placed in a rectangular fish tank (30 × 16 × 20 cm^3^) and recorded with a video camera for 20 minutes. Grooming behaviour was assessed from the last 10 minutes. This was to ensure the mice were habituated to their environment and minimise the effects of stress on grooming behaviour. A trained blinded observer manually scored and time-stamped grooming behaviour via behavioural observation research interactive software (BORIS) (Friard & Gamba, 2016). Percentage grooming time was calculated as the sum of time spent grooming or scratching the face, body, genitals, and tail, divided by the total duration (600 seconds) multiplied by 100. A grooming bout was recorded if the mouse groomed for greater than 4 seconds (Wilson et al., 2024). An interrupted bout was calculated when a mouse had stopped grooming for a period of < 4s, and then continued grooming.

### Statistical Analyses

Statistical analyses were performed using the GraphPad Prism 10.1.1 software package. Statistical significance was set at *p*□<□.05 for all experimental results. Multivariate ANOVAs were used according to the number of independent variables analysed. Repeated-measures ANOVA was adopted when there were multiple time points. In instances where Mauchly’s test revealed the assumption of sphericity to be violated in repeated-measures analysis, a Greenhouse-Geisser correction was applied. Post-hoc analysis was completed using the Holm-Šidák method or an unpaired t-test. When assessing the association between two variables, a parametric linear regression was used. In every behavioural experiment sex was considered an important biological variable. Thus, during our repeated measures LDB and grooming data we conducted individual three-way ANOVAs at each time-point to assess the effects of sex on these behaviours. Although, we found no significant sex differences in locomotor, HTR and the LDB test we did see significant sex differences at day 1 and 8 in the grooming behavioural test (Supplementary figures 1a and 1c), thus we decided to present male and female data separately.

## Results

### Psilocybin Increases Locomotion in WT but not in SAPAP3 KO mice

A repeated measures three-way ANOVA was used to assess how psilocybin and saline administration affected locomotor activity in WT and SAPAP3 KO mice (Figure 1). At baseline (i.e., the 5 minutes before injection) male and female SAPAP3 KO mice displayed less locomotor activity than WT mice (F_1,85_ = 85.41; *p* < 0.0001, η^2^ = 0.469) (Figure 1a). There were no significant sex differences at baseline (F_1, 85_ = 3.177; *p* = 0.0783; η^2^ = 0.018).

**Figure 1.**
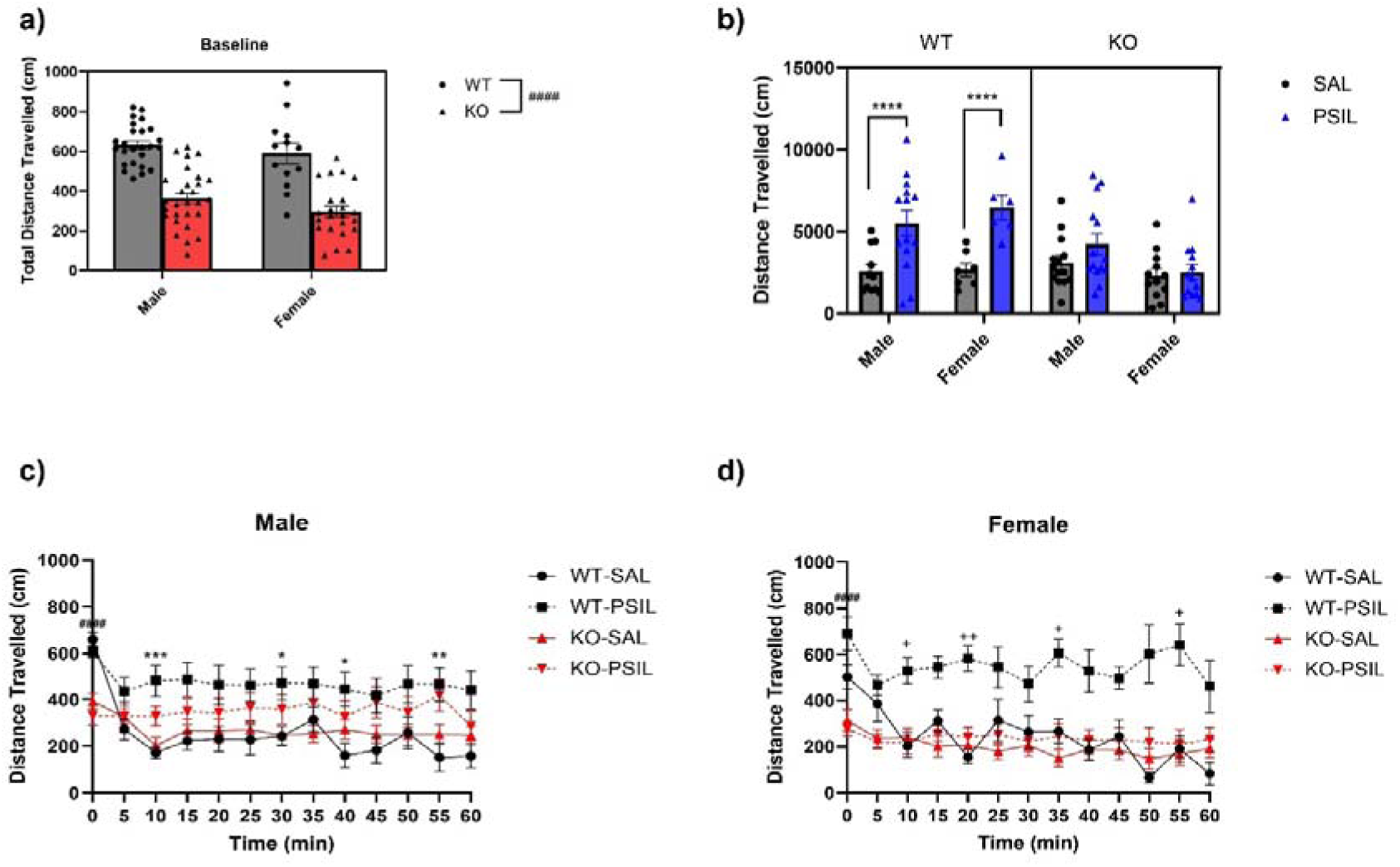
Locomotion Following Acute Psilocybin or Saline Administration **a)** SAPAP3 KO mice travelled less before drug administration (*p* < 0.0001). There were no significant differences between sexes (*p* = 0.078) and there was no significant interaction (*p* = 0.688). **b)** When analysing the cumulative distance travelled over the 60 minutes following drugs administration, we revealed a significant genotype x treatment interaction (*p* = 0.003) and found that psilocybin increased locomotion in WT mice (*p* < 0.0001) but not KO mice (*p* = 0.772). There was no significant sex effect (*p* = 0.401). No other significant interactions were observed. **c)** For male mice, a significant time x genotype effect (*p* < 0.0001), revealed that WT and KO mice had significantly different distance travelled in the 5 minutes before injection (time point 0) (*p* < 0.0001). Furthermore, a time x treatment effect revealed a significant locomotive difference between SAL and PSIL injected mice at 10 (*p* < 0.001), 30 (*p* = 0.048), 40 (*p* = 0.046) and 55 (*p* = 0.005) minutes after injection. *n* = 11 – 14 mice per group. **d)** For female mice, a significant time x genotype x treatment effect (*p* = 0.007) was observed. Post-hoc analysis also showed that at baseline (time point 0) WT mice had significantly different locomotion compared to KO mice (*p* < 0.0001). Additionally, it was found that WT mice receiving psilocybin had significantly different locomotion compared to WT mice receiving saline at 10 (*p* = 0.017), 20 (*p* = 0.002), 35 (*p* = 0.021) and 55 (*p* = 0.034) minutes after injection. No significant locomotive differences were found between SAPAP3 KO mice receiving SAL or PSIL at any time point (*p >* 0.930). *n* = 6 - 11 mice per group. WT = wild-type mice, KO = SAPAP3 knockout mice, SAL = saline, PSIL = psilocybin. ^####^*p* < .0001 genotype effect, \**p* < 0.05, \*\**p* < 0.01, \*\*\**p* < 0.001, \*\*\*\**p* < 0.0001 treatment effect, ^+^*p* < 0.05, ^++^*p* < 0.01 WT-PSIL vs WT-SAL.

For the total distance travelled in the 60 minutes following injection, a genotype x treatment interaction (F_1,83_ = 9.502; *p* = 0.003, η^2^= 0.076) revealed that psilocybin increased the distance travelled for WT (*p* < 0.0001) but not KO (*p* = 0.772) mice compared to saline (Figure 1b). There were no significant locomotor differences between male and female mice (F_1,83_ = 0.712; *p* = 0.401, η^2^ = 0.005).

When analysing how psilocybin altered locomotion across time in the first 60 minutes following administration, a time x treatment interaction was observed in males (F_12, 588_ = 4.37; *p* < .0001, η^2^ = .022). Šidák post-hoc analysis revealed that psilocybin increased locomotion in both WT and KO animals compared to saline treatment at 10 (*p* < .001), 30 (*p* = .048), 40 (*p* = .046) and 55 (*p* = .005) minutes after injection (Figure 1c). In females, a significant time x genotype x treatment effect occurred (F_12, 384_ = 2.32; *p* = .007, η^2^ = .016). Sidak multiple comparison analysis showed that while female WT mice treated with psilocybin had greater locomotion compared to WT mice treated with saline at 10 (*p* = .017), 20 (*p* = .002), 35 (*p* = .021) and 55 (*p* = .034) minutes after injection (Figure 1d), psilocybin did not significantly alter locomotion in female KO mice at any time point.

### Psilocybin Induces Head Twitches in WT and KO mice

A repeated measures three-way ANOVA was used to assess head twitches in male and female mice in the first 15 minutes following psilocybin or saline administration (Figure 2). In males, while a genotype effect did not reach significance (F_1, 51_ = 3.84; *p* = 0.054, η^2^ = 0.027), we found a significant time x treatment interaction (F_2,102_ = 8.16; *p* < 0.001, η^2^ = .027). Šidák post-hoc analysis revealed that psilocybin elicited a greater number of head twitches in mice at at 5 (*p* < .001) 10 (*p* < 0.0001) and 15 (*p* < 0.0001) minutes after injection compared to saline (Figure 2a). Similarly, in female mice there was no significant genotype effect (F_1,32_ = 3.13; *p* = 0.086, η^2^ = .044) but a significant time x treatment interaction (F_2,64_ = 3.80; *p* = 0.028, η^2^ = 0.018) was found.

**Figure 2.**
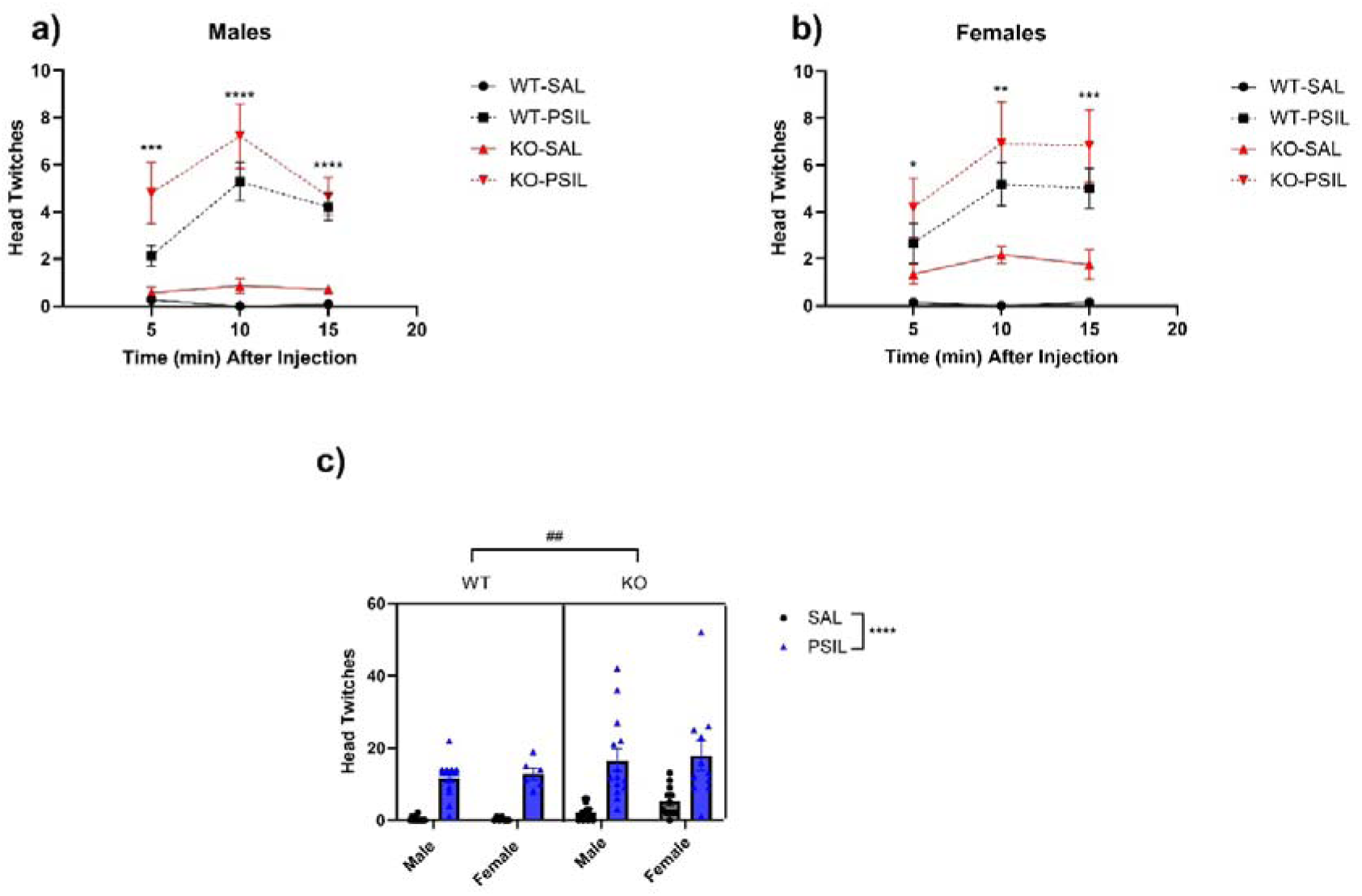
Head-Twitch Responses Immediately Following Psilocybin or Saline Administration **a)** A time x treatment interaction (*p* < 0.001) revealed a significant treatment effect at 5 (*p* < 0.001) 10 (*p* < 0.0001) and 15 (*p* < 0.0001) minutes after injection. No genotype effect was present (*p* = 0.055). *n* = 12 – 15 mice per group. **b)** A time x treatment interaction (*p* = 0.027) revealed a significant treatment effect at 5 (*p* = 0.021) 10 (*p* = 0.003) and 15 (*p* < 0.001) minutes after injection. A non-significant trend was observed for a genotype differences (*p* = 0.086). *n* = 6 – 12 mice per group. **c)** There were no significant differences in head twitches between sexes (*p* = 0.381). Psilocybin significantly increased the number of head twitches compared to saline (*p* < 0.0001). SAPAP3 KO animals had a greater number of head twitches compared to WT animals (*p* = 0.009). There were no significant interactions. WT = wild-type mice, KO = SAPAP3 knockout mice, SAL = saline, PSIL = psilocybin. \**p* < 0.05, \*\**p* < 0.01, \*\*\**p* < 0.001, \*\*\*\**p* < 0.0001 treatment effect. ^##^*p* < 0.01 genotype effect.

Psilocybin compared to saline, significantly increased the number of head twitches in both WT and KO animals 5 (*p* = 0.021), 10 (*p* = 0.003) and 15 (*p* < 0.001) minutes after injection (Figure 2b). We analysed the total head-twitch responses for the whole 15-minute duration and found that SAPAP3 KO mice had a significantly greater number of head twitches compared to WT mice (F_1,83_ = 7.168; *p* = 0.009, η^2^= 0.044). No significant sex differences were revealed (F_1,83_ = 0.776; *p* = 0.381, η^2^ = 0.005) (Figure 2c).

### Psilocybin Does Not Alter Anxiety-like Behaviour

A repeated measures three-way ANOVA was used to assess how psilocybin administration affected light duration in the LDB (Figure 3). In males, KO mice spent less time in the light compartment compared to WT mice (F_1,50_ = 12.38; *p* < 0.001, η^2^= 0.114) (Figure 3a). However, psilocybin administration did not alter light duration (F_1,50_ = 1.47; *p* = 0.231, η^2^ = 0.014) and light duration did not vary over time (F_2, 100_ = 0.55; *p* = 0.578, η^2^ = 0.004). There were no significant interactions. In females, a similar pattern of results occurred. KO animals spent less time in the light compartment compared to WT mice (F_1,32_ = 15.75; *p* < 0.001, η^2^ = 0.218) and psilocybin did not significantly alter light duration (F_1,32_ = 0.77; *p* = 0.385, η^2^ = 0.011) (Figure 3b). Light duration did not significantly vary over time (F_1.74, 55.80_ = 0.53; *p* = 0.565, η^2^ = 0.004). There were no significant interactions although a genotype x treatment effect was the closest to reaching significance (F_1, 32_ = 2.97; *p* = 0.097, η^2^ = 0.041).

**Figure 3.**
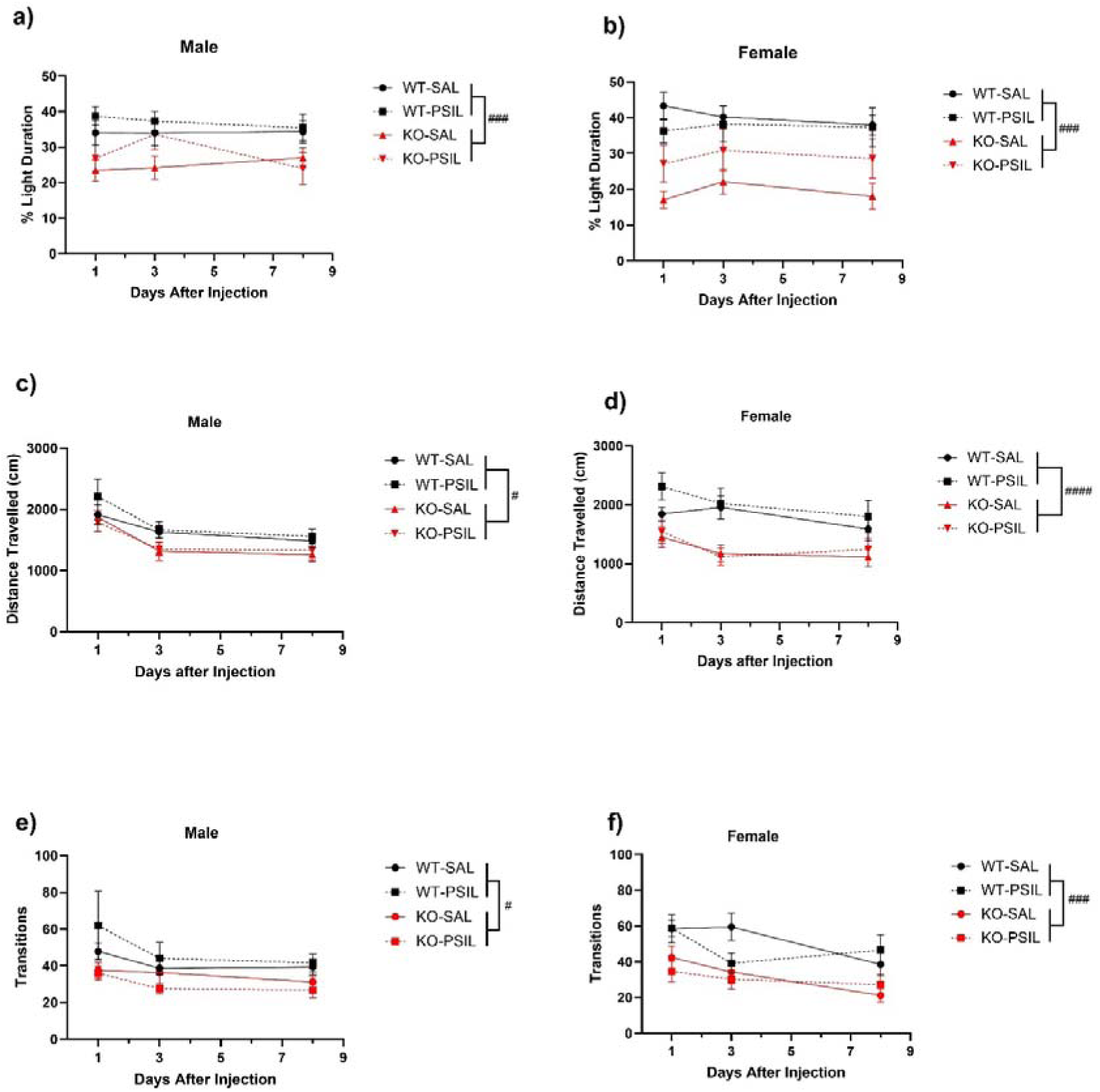
Effects of SAPAP3 Deletion and Psilocybin Treatment on Anxiety-Like Behaviour in the Light-Dark Box **a)** A main effect of genotype was present (*p* < 0.001), and there was no treatment or time effect (*p* = 0.231 and *p* = 0.578 respectively). *n* = 12 – 15 mice per group. **b)** A significant genotype effect was present (*p* < 0.001), and no treatment or time effect was observed (*p* = 0.385 and *p* = 0.565 respectively). There were no significant interactions (*p* > 0.05). **c)** Male SAPAP3 KO mice travelled significantly less than WT mice (*p* = 0.022). Time significantly affected distance travelled (*p* < 0.0001), however, psilocybin did not significantly alter locomotion (*p* = 0.524). No significant interactions occurred (*p* = 0.542). **d)** Similarly, female SAPAP3 KO mice travelled less than WT mice (*p* < 0.0001) and distance travelled significantly changed over time (*p* = 0.013). There was no significant effect of treatment (*p* = 0.300). No significant interactions occurred. **e)** Male SAPAP3 KO mice had less transitions than male WT mice (*p* = 0.012), but psilocybin (*p* = 0.822) or time (*p* = 0.089) did not alter transition behaviour. No significant interactions were present. **f)** for the females, KO animals also had less transition behaviour compared to WT mice (*p* < 0.001). Additionally, there was a main effect of time but not treatment (*p* = 0.433). No significant interactions were present, however a time x treatment interaction (*p* = 0.059) nearly reached significance. Males: *n* = 12 – 15 mice per group, Females: *n* = 6 – 12 mice per group. WT = wild-type mice, KO = SAPAP3 knockout mice, SAL = saline, PSIL = psilocybin. ^#^*p* < 0.05 genotype effect, ^###^*p* < 0.001, ^####^*p* < 0.0001 genotype effect.

Ancillary LDB measures revealed that male and female SAPAP3 KO mice travelled less than WT mice (F_1, 51_ = 5.62; *p* = 0.022, η^2^ = 0.042) (F_1,31_ = 20.19; *p* < 0.0001, η^2^ = 0.241) (Figure 3c and Figure 3d). There was no effect of treatment on distance travelled in the LDB in male (F_1,51_ = 0.41; *p* = 0.523, η^2^ = 0.003) and female mice (F_1,31_ = 1.11; *p* = 0.300, η^2^ = 0.013). Finally, KO animals had fewer transitions than WT animals (males: (F_1,51_ = 6.88; *p* = 0.012, η^2^ = 0.051), females: (F_1,32_ = 15.75; *p* < 0.001, η^2^ = 0.174) and psilocybin did not alter transition behaviour (Figure 3e and Figure 3f).

### Psilocybin Reduces Compulsive-like Grooming Behaviour in SAPAP3 KO Mice

A three-way repeated measures ANOVA was used to assess how psilocybin administration altered grooming behaviour in WT and SAPAP3 KO mice (Figure 4). In males, SAPAP3 KO mice significantly groomed more than WT mice (F_1,51_ = 89.91; *p* < 0.0001, η^2^= .384). Furthermore, a significant time x treatment x genotype effect was observed (F_2, 102_ = 4.58; *p* = 0.012, η^2^ = .019) and post-hoc analysis revealed that KO mice that received psilocybin had reduced grooming duration compared to KO mice that received saline at 3 (*p* = 0.010) and 8 days (*p* = 0.016) after injection (Figure 4a). There was no significant treatment effect on grooming for WT mice (*p* = 0.799). In females, SAPAP3 KO mice displayed more grooming than WT mice (F_1,32_ = 12.50; *p* = 0.001, η^2^ = .154). Psilocybin reduced grooming in both KO and WT mice (F_1,32_ = 4.99; *p* = 0.033, η^2^ = 0.064) (Figure 4b). Grooming duration was not significantly altered over time and no significant interactions were observed. Altogether, KO mice had a greater number of grooming bouts than WT mice (males: (F_1,52_ = 11.32; *p* = 0.014) females: (F_1,32_ = 7.58; *p* = 0.010)); and psilocybin did not alter the number of grooming bouts (males: (F_1,52_ = 0.05; *p* = 0.830) females: (F_1,32_ = 2.27; *p* = 0.014)) (Figure 4c and Figure 4d). Finally, there was a significant negative correlation between the number of head twitches and grooming duration in male (R^2^ = .24, *p* = 0.007) but not female mice (R^2^ = .08, *p* = 0.170).

**Figure 4.**
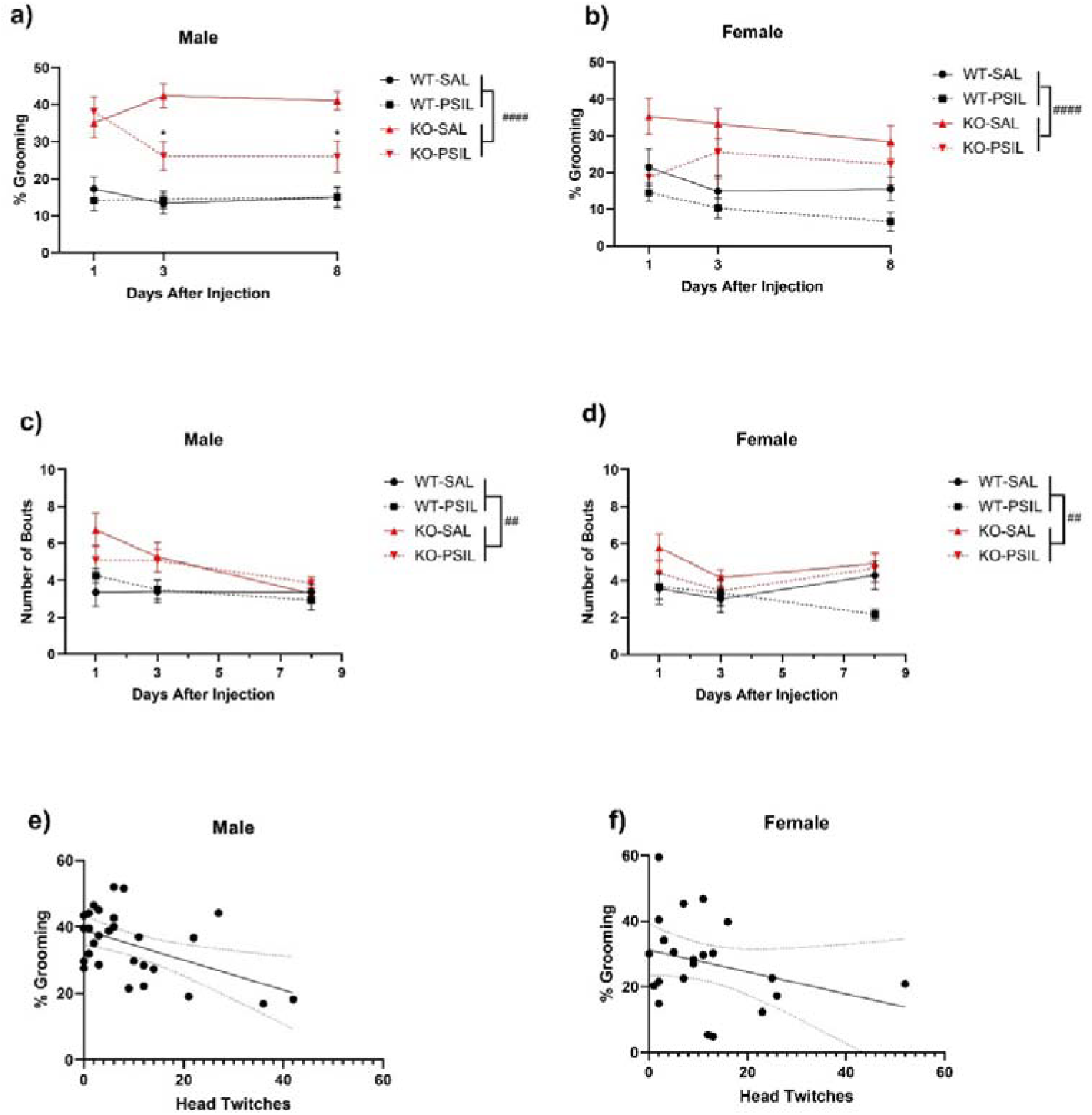
Effects of SAPAP3 Deletion and Psilocybin Treatment on Grooming Behaviour In male mice a significant genotype effect was observed (*p* < 0.0001) as well as a time x treatment x genotype effect (*p* = 0.013). Post-hoc analysis found a significant difference between KO-PSIL and KO-SAL mice at 3 (*p* = 0.010) and 8 days (*p* = 0.016) after injection. No significant differences were found between WT-SAL and WT-PSIL mice at any time-point (*p* = 0.844). *n* = 12 – 15 mice per group. **b)** For female mice, a significant main effect of genotype (*p* = 0.001) and treatment (*p* = 0.033) was found. No significant effect of time occurred (*p* = 0.358). There were no significant interactions. *n* = 6 – 12 mice per group. **c)** Male KO mice had a greater number of grooming bouts compared to WT mice (*p* = 0.001). Psilocybin did not alter the number of grooming bouts (*p* = 0.830). There was a significant effect of time (*p* = 0.002) and no significant interactions. **d)** There was a significant genotype (*p* = 0.010) but not time (*p* = 0.228) or treatment effect (*p* = 0.142) in female mice. No significant interactions occurred. **e)** There was a significant association between head twitches and grooming duration in male mice treated with psilocybin (*R*^2^ = 0.24, *p* = 0.007). **f)** there was no significant association between head twitches and grooming in female mice treated with psilocybin (*R*^2^ = .08, *p* = 0.149). Males: *n* = 12 - 15 mice per group. Female: *n* = 6 – 12 mice per group. WT = wild-type mice, KO = SAPAP3 knockout mice, SAL = saline, PSIL = psilocybin. ^##^*p* < 0.01, ^###^*p* < 0.001, genotype effect. \**p* < 0.05, \*\**p* < 0.01 KO-SAL vs KO-PSIL. <u>Note:</u> The grooming values were the mean grooming scores for each mouse across all time 3 points. *n* = 28 mice for males and *n* = 23 mice for females. Additionally, we only included KO-PSIL mice to avoid the confounds of treatment and genotype effects.

To assess sex effects on grooming, we ran a three-way ANOVA at each time-point. We found that one day after psilocybin administration there was a significant treatment x sex interaction (F_1,83_ = 4.08; *p* = 0.048, η^2^ = 0.03) and post-hoc analysis revealed that psilocybin reduced grooming duration compared to vehicle in female (*p* = 0.032) but not male (*p* = 0.955) mice (Supplementary figure 1a). Three days after psilocybin administration male and female mice did not have significantly different grooming (F_1,83_ = 1.01; *p* = 0.317, η^2^ = 0.007) (Supplementary figure 1b). Finally, 8 days post drug administration, male mice had a significantly greater grooming duration compared to female mice (F_1,83_ = 4.77; *p* = 0.032, η^2^ = 0.03) (Supplementary figure 1c).

## Discussion

This study is the first to investigate the effect of psilocybin administration in the SAPAP3 KO mouse model, which is a well-validated mouse model of compulsive-like behaviours. As expected, psilocybin robustly increased head-twitches at the dose used (1 mg/kg), indicative of its hallucinogenic-like properties. Notably, SAPAP3 KO mice exhibited a greater number of head-twitches compared to WT mice. Furthermore, we found that psilocybin increased locomotion in WT mice but not in SAPAP3 KO mice. Although psilocybin did not alter anxiety-like behaviour, we found that a single dose of psilocybin reduced compulsive-like grooming behaviour. Notably, this effect was long-lasting (i.e. occurring at 3- and 8-days post injection) and mainly observed in the SAPAP3 KO mice. These novel findings provide new insights into the effects of psilocybin on psychiatric endophenotypes and support the treatment potential of psilocybin for patients with OCRD.

### Validating the Anxiety- and Compulsive-like Behaviours of SAPAP3 KO mice

In alignment with previous literature (Ade et al., 2016; Gattuso et al., 2023a; Welch et al., 2007), we found that SAPAP3 KO mice exhibited increased anxiety-like behaviour by spending less time in the light compartment of the LDB compared to WT mice. We also found that SAPAP3 KO animals had fewer transitions in the LDB compared to WT animals which is indicative of reduced exploratory behaviour. Furthermore, as expected SAPAP3 KO mice groomed significantly more than WT mice (Davis et al., 2021; Gattuso et al., 2023a; Lamothe et al., 2023; Ramírez-Armenta et al., 2022; Wan et al., 2014; Welch et al., 2007). Finally, we and others (Hadjas et al., 2020) have previously found that SAPAP3 KO mice display a hypolocomotion phenotype (Gattuso et al., 2023a), which was also replicated in this study.

### Psilocybin acute behavioural responses

#### Locomotor activity

The acute behavioural effects of psilocybin remain unclear as studies have typically used a variety of doses and different animal models. Acute injections at higher doses of psilocybin, such as 3 and 5 mg/kg, have been known to reduce locomotor activity in mice (Fadahunsi et al., 2022; Halberstadt & Geyer, 2011; Hesselgrave et al., 2021; Rahbarnia et al., 2023). At the same dose (1 mg/kg) as our study, but in a different mouse strain, Odland et al. (Odland et al., 2021) found that psilocybin did not alter locomotion in female NMRI mice. Over a 24-hour period Alper et al. (2023) found that a single dose of psilocybin (1 mg/kg), immediately before the dark cycle did not alter distance travelled in male and female C57BL/6J mice. However, they did find that (at a dose of 0.5 mg/kg) psilocybin increased locomotion in female mice for the first 8 hours. Similarly, Rahbarnia and colleagues (2023) found that psilocybin (1 mg/kg, i.p.) did not alter locomotion in male C57BL/6J mice.

In the present study, we found that WT mice had greater locomotor activity in the 60 minutes following acute psilocybin i.p. injection (1 mg/kg). These effects seem to be driven by 5-HT_2A_ receptor agonism, as it has been found that the hallucinogen and 5-HT_2A_ receptor agonist 1-(2,5-dimethoxy-4-iodophenyl)-2-aminopropane (DOI) increases locomotion at lower doses (0.625 – 5.0 mg/kg) and decreased locomotor activity at higher doses (≥10□mg/kg) (Halberstadt et al., 2009). Furthermore, Halberstadt et al. (2009) found that DOI at 1 mg/kg increased locomotion in C57BL6/J mice; however, did not alter locomotion in 5-HT_2A_ receptor KO mice. However, when DOI reduces locomotion at higher doses, this effect seems to be mediated by the 5-HT_2C_ receptor, as the selective 5-HT_2C_ antagonist attenuated the decreases in locomotion induced by DOI (10 mg/kg) (Halberstadt et al., 2009). Furthermore, the classical psychedelic, lysergic acid diethylamide (LSD) also seems to induce 5-HT_2A_ receptor mediated hyperlocomotion (Rodriguiz et al., 2021). Additionally, 5-HT_2A_ receptor antagonism leads to decreased locomotion in C57BL6/J mice (Rahbarnia et al., 2023). Thus, our data adds credence to the notion that psilocybin at higher doses may decrease locomotion via a greater agonism of the 5-HT_2C_ receptor, whereas lower doses such as 1 mg/kg may increase locomotion via 5-HT_2A_ mediated effects. However, such findings are mixed (Rahbarnia et al., 2023) and this would need to be confirmed by future molecular experiments.

Interestingly, we also observed that psilocybin did not increase locomotion in SAPAP3 KO mice compared to saline. If the increased locomotion is driven by 5-HT_2A_ receptor activity, our observed results may suggest altered *in vivo* 5-HT_2A_ receptor function in SAPAP3 KO mice. Supporting this, we also found that SAPAP3 KO mice had altered HTR compared to WT littermate controls, and the HTR is a 5-HT_2A_ receptor mediated behaviour (Halberstadt et al., 2020; Jaster et al., 2022; Shahar et al., 2022). However, at this stage, this particular evidence is exploratory and could be confirmed by subsequent causal behavioural pharmacological experiments.

#### Head-Twitch Response

The head-twitch response (HTR) in mice is a proxy for the hallucinogenic experience in humans and is driven by 5-HT_2A_ receptor agonism (Halberstadt et al., 2020). At the dosage used in our study, we found that psilocybin substantially increased HTR in WT and SAPAP3 KO mice compared to saline administration. These findings in WT mice are consistent with previous literature (Hesselgrave et al., 2021). Additionally, we found a significant effect of genotype in the HTR, where SAPAP3 KO mice displayed a greater number of head twitches compared to WT mice. These results are likely due to the tic-like body/head shakes recently observed in SAPAP3 KO mice (Lamothe et al., 2023). In contrast to our locomotion data, the lack of a genotype x treatment interaction suggests that although psilocybin significantly increased the number of head twitches compared to saline, there was no significant difference between WT and SAPAP3 KO mice.

### Effects of Psilocybin on Anxiety- and Compulsive-Like Behaviour

Preclinical investigations into psilocybin on anxiety-like behaviour exhibit large heterogeneity, with studies finding mixed results, assessing behaviour in various behavioural assays, using different animal models and dosing paradigms. For instance, Hibicke et al. (2020) found that a single dose of psilocybin (1 mg/kg i.p.) led to enduring reductions (41 days post psilocybin administration) in anxiety-like behaviour in the elevated-plus maze (EPM) of male Wistar-Kyoto rats. Jones et al. (2023) found that 15 minutes after injection psilocybin (3 mg/kg, i.p., single dose) increased anxiety-like behaviour in male C57Bl6/J mice by reducing time spent in the middle of the open-field test (OFT), but 4 hours after injection, mice displayed anxiolytic behaviour by spending more time in the centre of the OFT. The anxiolytic effect was also found 7 days after psilocybin administration.

Finally, Wojtas et al. (2022) found that neither a single dose of psilocybin at 2 mg/kg or 10 mg/kg (i.p.) altered anxiety-like behaviour in the OFT and the LDB 24 hours after administration in male Wistar Han Rats. Thus, our study was the first to assess how psilocybin affects anxiety-like behaviour in the LDB for mice. We found that psilocybin did not alter anxiety-like behaviour in the LDB for WT or KO animals.

However, a single dose of psilocybin (1 mg/kg, i.p.) led to enduring reductions in compulsive-like grooming behaviour in male SAPAP3 KO mice and reduced grooming behaviour in both SAPAP3 KO and WT female mice. In male SAPAP3 KO mice, psilocybin significantly reduced grooming behaviour at 3 and 8 days after injection compared to saline-treated male KO mice. Psilocybin did not alter grooming in male WT mice; however, in the female mice, psilocybin reduced grooming mice across all time points (1, 3 and 8 days after injection) in both genotypes. Interestingly, we found that there was a significant negative association between grooming duration and the number of head twitches in male KO mice. Thus, on average, male KO mice that tended to have a greater number of head twitches also had a greater reduction grooming behaviour. Although, these findings are correlational and we cannot imply causation, they do suggest that the acute head twitch behavioural response may be necessary for the reduction in grooming behaviour in male KO mice. However, it is important to consider that this effect was not seen in female KO mice which may be because we saw an overall treatment effect with the female mice rather than a genotype x treatment effect. Our findings suggest that blockade of the HTR in male mice might abolish the therapeutic effects of psilocybin observed in this study but this would need to be confirmed by future behavioural pharmacological studies.

As we observed behavioural changes at 3- and 8-day time points after injection, but no acute changes at 1-day post injection in male SAPAP3 KO mice, this could be due to a transient alteration in serotonergic neurotransmission. For instance, upon administration, serotonergic psychedelics have been found to acutely internalise 5-HT_2A_ receptors with re-expression occurring up to 48 hours later (Darmon et al., 2015; Nichols, 2016). However, this possible explanation would only be applicable to the male KO mice as psilocybin reduced grooming behaviour in female mice across all time points. As psilocybin’s reduction in grooming behaviour occurred in both genotypes in females, this effect might not reflect reduction in compulsive grooming *per se*, but may rather be indicative of increasing stress-resiliency, which has been previously characterised with psilocybin (Hibicke et al., 2023; Kiilerich et al., 2023). Indeed, grooming behaviour is modulated by stress in mice (Smolinsky et al., 2009). However, psilocybin did not alter the number of interrupted bouts (data not shown) in WT or KO mice which is sensitive to stress effects (Kalueff et al., 2016). Thus, future research could investigate how psilocybin alters additional compulsive behavioural assays such as reversal learning (a measure of behavioural flexibility related to compulsive behaviour (Robbins et al., 2024) and found to be impaired in SAPAP3 KO mice (Manning et al., 2019; van den Boom et al., 2019)) and schedule-induced polydipsia (Robbins et al., 2024). Testing how psilocybin affects these additional compulsive-like behaviour will enable researchers to further test whether the reduction in grooming behaviour in female mice is due to a reduction on compulsive-like behaviour.

Currently, some research suggests that psilocybin’s therapeutic effect is mediated by the 5-HT_2A_ receptor (Catlow et al., 2013; Hagsäter et al., 2021; Meinhardt et al., 2021) whereas other research provide support for 5-HT_2A_ receptor independent effects (Hesselgrave et al., 2021; Moliner et al., 2023; Shao et al., 2021; Yaden & Griffiths, 2020). In the context of compulsive-like behaviour, psilocybin (2mg/kg i.p.) reduced marble burying in female NMRI mice and the effect was independent of the 5-HT_2A_ and 5-HT_2C_ receptor (Odland et al., 2021). Additionally, psilocybin (4.4 mg/kg, i.p.) reduced marble burying in male ICR mice and the effect was not attenuated by the 5-HT_2A_ antagonist M100907 or 5-HT_1A_ antagonist WAY100635. In addition to the 5-HT_2A_, 5-HT_1A_ and 5-HT_2C_ receptor psilocybin also has high affinity for the serotonin transporter and the TAAR1 receptors (Rickli et al., 2016) and it has been argued that these receptors may also contribute to the anti-compulsive-like effects of psilocybin and could be within the purview of future research (Singh et al., 2023). A limitation of our study is that we did not pretreat the SAPAP3 KO mice with any 5-HT (or dopaminergic) receptor antagonists so we cannot decipher which receptors may be necessary for the reduced grooming behaviour observed in this study. Furthermore, to further understand the neural underpinnings of the psilocybin-induced reduction in grooming behaviour, C-fos expression following grooming behaviour in psilocybin-treated mice could be an exciting future direction.

These sexually dimorphic effects of psilocybin (evidenced by our sex x treatment interaction at day 1 of grooming – Supplementary Figure 1a) on grooming behaviour are novel and interesting. They could be due to many possible factors, including genetic differences which lead to sex-specific drug metabolism (Mogil & Chanda, 2005), hormonal variations such as estrogen which can affect enzymes related to drug metabolism (Gandhi et al., 2004), and immune system variation, with females having a stronger immune response compared to males which can influence the efficacy of drugs (such as psilocybin (Mason et al., 2023)) which alter the immune system (Klein & Flanagan, 2016).

Our findings greatly extend—the preclinical literature assessing psilocybin administration on compulsive behaviour (Matsushima et al., 2009; Odland et al., 2021; Singh et al., 2023). As SAPAP3 KO mice are a preclinical model of compulsive behaviour relevant to a range of clinical populations across the OCRD spectrum such as OCD, trichotillomania, Tourette syndrome and excoriation disorder (Lamothe et al., 2023) our findings support the research into psilocybin treatment for these clinical groups. Due to the enduring therapeutic effects of a single dose of psilocybin and the fact that psilocin has a plasma clearance time of about 4 hours in mice (Jones et al., 2023), we hypothesise that psilocybin has induced structural alterations in the brain. Indeed, psilocybin has been found to increase spinogenesis (Moliner et al., 2023; Shao et al., 2021), neuronal proliferation, differentiation survival and maturation (Catlow et al., 2013) and upregulate neuroplasticity-related gene expression (Davoudian et al., 2023; Fadahunsi et al., 2022). However, assessing the cellular mechanisms by which psilocybin reduced compulsive behaviour was beyond the scope of this study, but could be within the purview of future research. Additionally, as grooming behaviour was still reduced at 8 days following injection and this was our last time point, it is unknown how long these therapeutic effects might last. Findings in rats have revealed therapeutic effects lasting as long as 6 weeks from a single dose of psilocybin (Hibicke et al., 2020). Interestingly, the study found that psilocybin led to persisting antidepressant-like effects whereas ketamine did not. Similarly, our findings have found persisting reductions in compulsive-like behaviour (particularly in male KO mice), whereas ketamine has not (Davis et al., 2021; Gattuso et al., 2023a). In humans, sustained therapeutic effects from 1-3 doses of psilocybin have been reported for as long as months and even years after administration (Agin-Liebes et al., 2020; Gukasyan et al., 2022; Johnson et al., 2017) which align with our findings and the findings of (Dutta & Sengupta, 2016) who calculated that 8 adult mouse days is approximately equal to 2 adult human years.

## Conclusion

Our study is the first to assess the effects of psilocybin administration in the SAPAP3 KO mouse model, a well-validated mouse model of compulsive behaviour and relevant to individuals diagnosed with OCRD (Lamothe et al., 2023). We confirmed that SAPAP3 KO mice exhibited increased anxiety-like and compulsive-like behaviour. Our study has revealed novel behavioural pharmacology findings and included sex as a factor which has been inadequately analysed in previous studies (Pedicini & Cordner, 2023). We found that the increased locomotion following acute administration of psilocybin was only observed in WT but not in KO mice. Furthermore, the dose used was capable of eliciting a substantial increase in the number of head twitches for both WT and KO animals, indicating its hallucinogenic properties at the dose used. We found that although a single dose of psilocybin did not alter anxiety-like behaviour it did lead to enduring reductions in grooming behaviour in SAPAP3 KO mice. Specifically, psilocybin decreased compulsive-like grooming behaviour at 3 and 8 days after injection, in male KO mice but not WT mice, however, decreased grooming behaviour in female WT and KO mice at 1, 3 and 8 days after injection. These sex differences may suggest that psilocybin’s therapeutic response may differ between sexes. As current frontline treatments for OCD (e.g. SSRIs such as fluoxetine), only reduced compulsive-like grooming behaviour after 6 days of chronic administration, but not after a single injection (Welch et al., 2007) our current study has revealed that psilocybin is a prime candidate molecule for the treatment of OCD and OCRD. However, we did not directly compare the effects of fluoxetine to psilocybin under the same experimental conditions, something which could be within the scope of future investigations. Future studies could investigate whether the anti-compulsive-like effects of psilocybin reported here are specific to psilocybin or extend to other psychedelic compounds such as LSD and N,N-Dimethyltryptamine (DMT) and non-hallucinogenic psychedelic analogues. Finally, it would be interesting to compare our results with other doses of psilocybin. For instance, would a higher dose of psilocybin (i.e., 3 mg/kg - 5mg/kg) lead to both a reduction in anxiety-like and compulsive-like behaviour? Furthermore, would a low dose of psilocybin (i.e., 0.05 – 0.1 mg/kg) that did not significantly invoke head twitches still lead to the reduction in grooming behaviour observed in the current study?

## Acknowledgements

We thank Prof Susanne Ahmari and Dr Elizabeth Manning for generously providing breeding pairs of SAPAP3 heterozygous mice and helping us setting up SAPAP3 KO mice colony at the Florey Institute. We acknowledge Shan Li (Research Assistant in the Hannan Lab) and Florey Core Animal Services (especially Brett Purcell who runs the Florey Behaviour Facility). We acknowledge Usona Institute Inc. which provided psilocybin for that study. The Florey Institute of Neuroscience and Mental Health acknowledges the support from the Victorian Government’s Operational Infrastructure Support Grant.

