## Supplementary File for "Psilocybin Reduces Grooming in the SAPAP3 Knockout Mouse Model of Compulsive Behaviour"

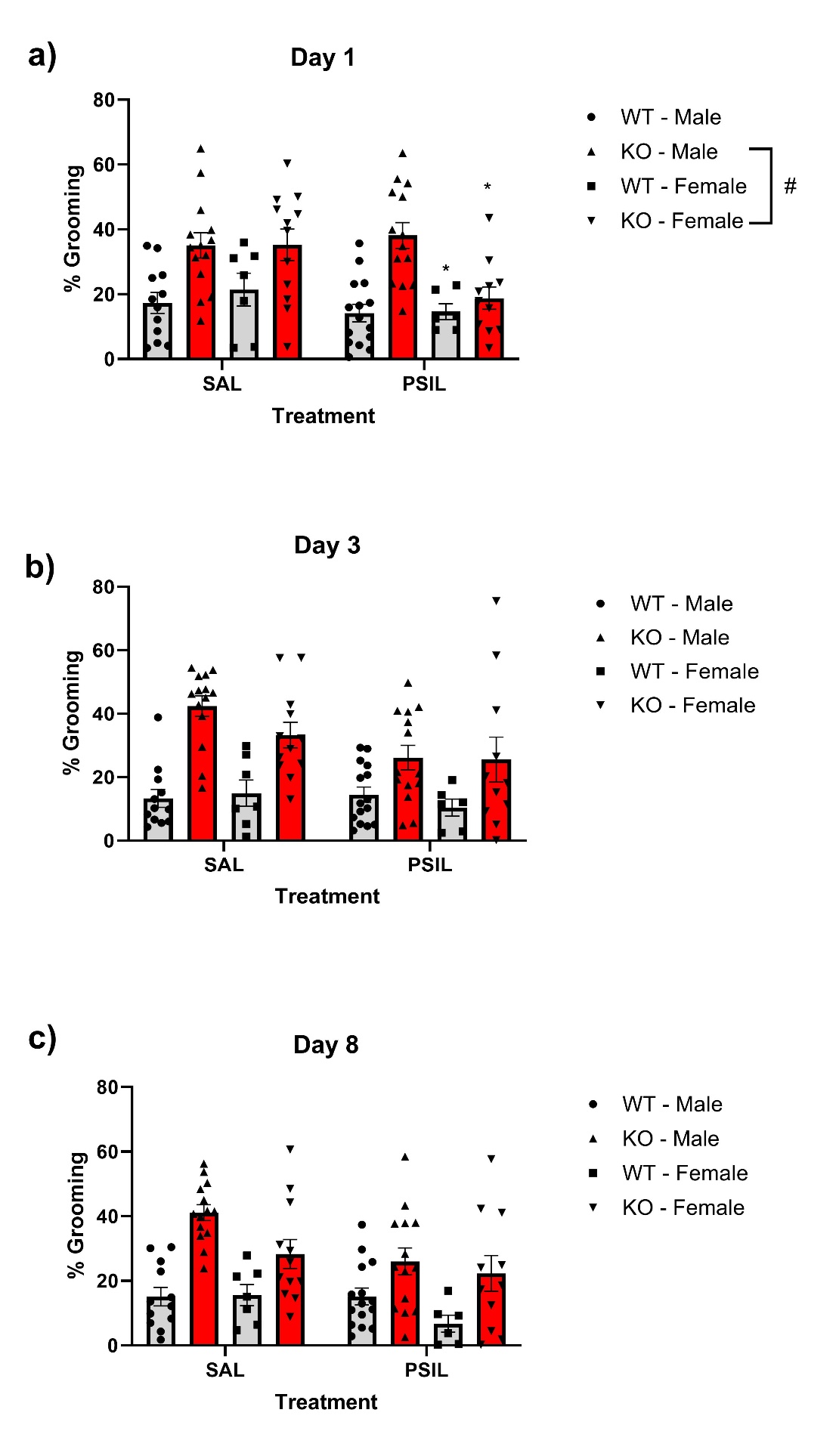

**Supplementary Figure 1. Effects of SAPAP3 Deletion, Psilocybin Treatment and Sex on Grooming Behaviour**

**a)** One day post psilocybin administration, SAPAP3 KO mice groomed significantly more than WT mice (*p* < 0.0001). Additionally, a sex x treatment interaction was observed (*p* = 0.481) and post-hoc analysis revealed that psilocybin significantly reduced grooming in female (*p* = 0.032) but not male mice (*p* = 0.955). Additionally, a sex x genotype effect (*p* = 0.043) revealed that male KO mice groomed significantly more than female KO mice (*p* = 0.035). **b)** Three days post psilocybin administration, there was a main effect of treatment (*p* = 0.026) and genotype (*p* < 0.0001). However, there were no sex differences (*p* = 0.317) or significant interactions. **C)** Eight days post psilocybin administration, there was a main effect of treatment (*p* = 0.009), sex (*p* = 0.032) and genotype (*p* < 0.0001).
